# groov^DB^ in 2025: A community-editable database of small molecule biosensors

**DOI:** 10.1101/2025.08.18.670880

**Authors:** Joshua D. Love, Brady M. Rafferty, Michael Thomas, Nicole Zhao, Pranay Talla, Michael Springer, Pamela A. Silver, Simon d’Oelsnitz

## Abstract

The groov^DB^ database (https://groov.bio) was launched in 2022 with the goal of organizing information on prokaryotic ligand-inducible transcription factors (TFs). This class of proteins is important in fundamental areas of microbiology research and for biotechnological applications that develop biosensors for diagnostics, enzyme screening, and real-time metabolite tracking. Uniquely, groov^DB^ contains stringently curated, literature-referenced data on both TF:DNA and TF:ligand interactions. Here, we describe a major technical update to groov^DB^, making the database community-editable and adding several advanced features. Users can now add new TF entries and update existing entries using a simple online form. New user interface elements display interactive protein structures and DNA-binding motifs. Updated query methods enable database searches via text, chemical similarity, and attribute-filtering. A new data architecture reduces page load time by five-fold. Finally, the number of TF entries has more than doubled and all source code is now open-access.

**Graphic abstract:** 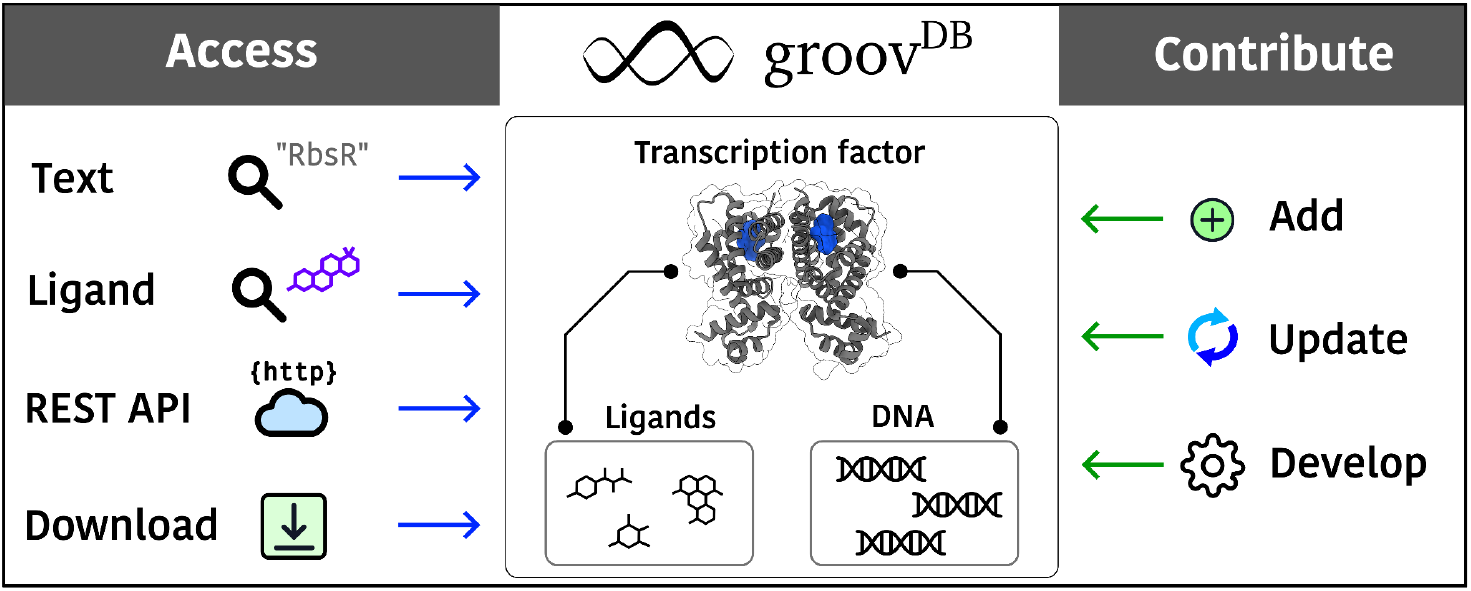

## Introduction

Prokaryotes adapt to their environment by triggering the expression of genes in response to chemical stimuli using ligand-inducible transcription factors (TFs). Upon binding to their cognate small-molecule ligand, transcription factors undergo a conformational change that alters their DNA-binding affinity. These structural shifts can either facilitate transcription, by recruiting RNA polymerase to the promoter, or inhibit transcription, by obstructing polymerase access to the initiation site^1^. TFs largely define how a prokaryote’s chemical environment influences its transcriptional program, and therefore mapping the ligand and DNA targets of TFs is crucial to understand and predict microbial behavior.

For decades, prokaryotic TFs have also played a pivotal role in biotechnology. Initially employed as inducible gene expression systems in organisms such as yeasts, mammalian cells, insects, and plants, TFs became core to the field of Synthetic Biology in the early 2000s, where they were used to build logic circuits^2,3^. Shortly afterwards, TFs found use as high-throughput screening tools for strain and enzyme engineering; by controlling GFP expression using a TF responsive to an enzyme product, millions of enzyme variants expressed within cells can be screened using fluorescence-activated cell sorting^4–7^. More elaborate genetic schemes involved using TFs to stabilize production phenotypes within engineered strains, dynamically regulate pathways to minimize cellular toxicity, and autonomously induce gene expression based on cellular density to obviate the need for inducers^8–11^. Beyond metabolic engineering applications, TFs have served as central components for diagnostics and live cell imaging. Cell-free systems controlled by TFs have been used to monitor water pollution and build responsive textiles^12,13^. TFs have been linked to bioelectronic and acoustic readouts to report on disease states within the gut of live animals^14–16^. And TFs fused to split GFP proteins have been used to record real-time abundance of compounds such as hydrogen peroxide, cyclic di-GMP, itaconate, and fructose 1,6-bisphosphate in mammalian cells^17–20^.

Despite these exciting applications, identifying a ligand-inducible TF appropriate for a given project remains challenging. Few existing resources document experimentally validated TF binding interactions, and in particular, ligand binding interactions. Several databases, such as MIST, P2CS, KEGG, and iModulonDB, describe signal transduction pathways or transcriptomic datasets in prokaryotes, but lack specific DNA or ligand binding information^21–24^. Other databases, such as RegulonDB, DBTBS, CoryneRegNet, and RhizoRegNet, report on the DNA binding interactions of TFs, but information is only available for specific organisms^25–28^.

Computational tools exist for predicting candidate inducer molecules or DNA operators of TFs, such as TFBMiner, Snowprint, and Ligify, but lack experimental validation^29–31^. The SensiPath tool provides enzyme cascades for molecule detection, but pulls from outdated TF databases^32^. Some databases suggest candidate TF-ligand interactions, such as RegPrecise and PRODORIC, but these interactions are often not supported by experimental evidence^33,34^. Finally, an excellent dataset of “detectable molecules” has been manually curated from literature sources, but lacks detailed information on transcription factor identity, experiments conducted, and DNA binding information^35^.

In 2022, we released groov^DB^, a manually curated database of experimentally validated ligand and DNA interactions for prokaryotic transcription factors^31^. Over the past few years, groov^DB^ has been used to benchmark and augment computational tools for biosensor design^32,33^. In this release, new features and updated content are incorporated to make the database easier to query, more responsive, more intuitive, and open to community contribution. Overall, groov^DB^ is positioned to facilitate the identification of TFs broadly useful for microbiology and biotechnology research.

### System overview and database content

Since its launch, groov^DB^ has been a useful resource for ligand-inducible transcription factors, with 10-20 active users each day. In this latest technical update, we have made groov^DB^ a community-editable and open-source application. In addition to submissions from the groov^DB^ curation team, users can now add their own transcription factors or update existing entries directly using a simple online form. In addition, the open-sourced codebase for both the frontend and backend, alongside detailed documentation, now enables software contributions from the community. Together, these updates greatly facilitate both internal and external contributions to this growing database. Finding TF entries is now easier with the ability to perform advanced searches based on TF property filters, text-based queries, and a new chemical similarity search feature. We also offer new methods to access data within groov^DB^, including programmatic access via a REST API and downloading the entire database as a static file. Improved user interface elements support the display of interactive Alphafold structures when crystal structures are not available, as well as DNA motif visualization that summarizes interactions with multiple DNA sequences. Finally, we have more than doubled the size of groov^DB^, which now contains >215 unique transcription factors, >260 unique operator sequences, and >320 unique ligands from 112 unique organisms.

### Community editing

To facilitate the continuous expansion of content in groov^DB^, we have introduced a feature allowing users to add or update TF entry pages after creating an account. Users can create or sign into their account simply using their Google credentials, via the secure OAuth 2.0 protocol, or by manually entering their account details. User authentication and authorization is securely managed by the AWS Cognito service. After sign-in, users can create a TF entry by completing a form that includes necessary metadata, ligand binding information, and DNA binding information for their new entry. Criteria for ligand and DNA interaction evidence are thoroughly described in documentation pages, helping guide the contributor while maintaining the rigorous integrity of the database. Inputs are validated in real time to prevent downstream integration issues. Upon submission, the TF entry is moved to a queue that must be reviewed and approved by an administrator, which further strengthens the accuracy of the database content. After approval, a bioinformatic pipeline is run to fetch additional metadata, such as PDB structure codes, the Alphafold structure code, host organism name and taxonomy ID, the protein sequence, genome context, additional references, and links to the external databases such as KEGG^23^. Following a second round of administrator approval to validate the newly generated content, the TF entry is added to the public-facing database. Users can also update existing TF entries to ensure database content is current and up-to-date as new literature results are published. To update a TF entry, users edit the entry’s populated submission form. The updated TF form is then passed through the same administrator review process as for TF entry creation to ensure consistency. Users can also submit batches of sensors by contacting the groov^DB^ curators directly. Finally, beyond content contributions, users can now support the development of the open-source groov^DB^ software, which is freely available on GitHub as frontend and backend repositories.

### New query tools

groov^DB^ provides three methods to query the database: browsing, text search, and chemical similarity search. Users can browse for TFs based on several features, such as their structural family, their host organism, Uniprot ID, or RefSeq ID through a new interactive data table on the “Browse” page. Pages for structural families include a brief description of representative features for regulators within that family, such as mechanism and ligand structures. Advanced searches can be performed in the same data table by filtering entries based on TF attributes or via a comprehensive full text-based search. From the Home page, the database can be queried using text-based and chemical-similarity search. Text searches return entries that have TF aliases or ligands that match the text input. Chemical similarity search accepts an input chemical in SMILES notation and returns TF entries with corresponding ligands that are similar to the input chemical, based on Tanimoto similarity scores. During this process, the input SMILES code is converted into a Morgan fingerprint using RDKit (radius: 2, size: 2048), and compared to fingerprints for all ligands in groov^DB^. Only ligands that meet a similarity threshold are returned, and this threshold can be adjusted by the user to tune the stringency of the search. The chemical similarity search method is particularly useful when users are looking for a template TF biosensor for evolution that responds to a ligand similar to a user-defined target, as several directed evolution methodologies exist for altering TF ligand specificity^6,38–41^. As the database content continues to grow, these query features will greatly simplify finding relevant TF entries.

### Programmatic access

New methods are provided for programmatic access to all data in groov^DB^, which is particularly valuable for bioinformatics or machine learning projects that need high-quality, manually curated data. A new REST API allows users to fetch subsets of the database, including defined TF entries, sets of entries from particular families, or all entries. Detailed documentation on query structures with examples and the expected data format to be returned is provided in a dedicated Swagger page. The full database in JSON format is also available via a download link, which is continuously updated as TF entries are added or updated.

### Enhanced web interface

The web interface of groov^DB^ has been significantly revised to provide a more intuitive and data-rich user experience. Several improvements have been made to the TF entry pages (Figure 1). Interactive Alphafold structures fetched from the Alphafold Protein Structure Database are now supported for all TF entries, which aids the molecular exploration of DNA and ligand binding interfaces^42^. DNA motif visualization has also been improved. In the previous version, only one DNA sequence could be associated with each TF entry, which limited accuracy. The current release now supports the integration of multiple DNA sequences, which are used to create a position-weighed sequence motif using the React LogoJS component^43^. This change is particularly important for accurately reporting the DNA binding specificity of global TF regulators, which often promiscuously bind to many DNA sequences bearing a small specificity motif^1^. Additional display modifications further improve the readability of data in each TF entry page, including a new protein sequence viewer that indicates the protein length as well as new Uniprot-like display formats for metadata, such as the organism name and KEGG ID. On the Home page, a continuously updated panel with database content statistics, including the number of unique TFs and ligands, is now displayed. A new Tools tab links to useful resources for the design and discovery of TF biosensors, including Ligify, Snowprint, TFBMiner, and SensBio^29–31,37^. A toggleable dark mode has been added, and components within sensor entry pages can be viewed either as a single page or as tabs. Finally, documentation on the About page has been significantly revised. Criteria for accepted TF:ligand and TF:DNA interaction evidence is detailed, a new Frequently Asked Questions page is added, and a new contact form with CAPTCHA validation is incorporated.

**Figure 1:**
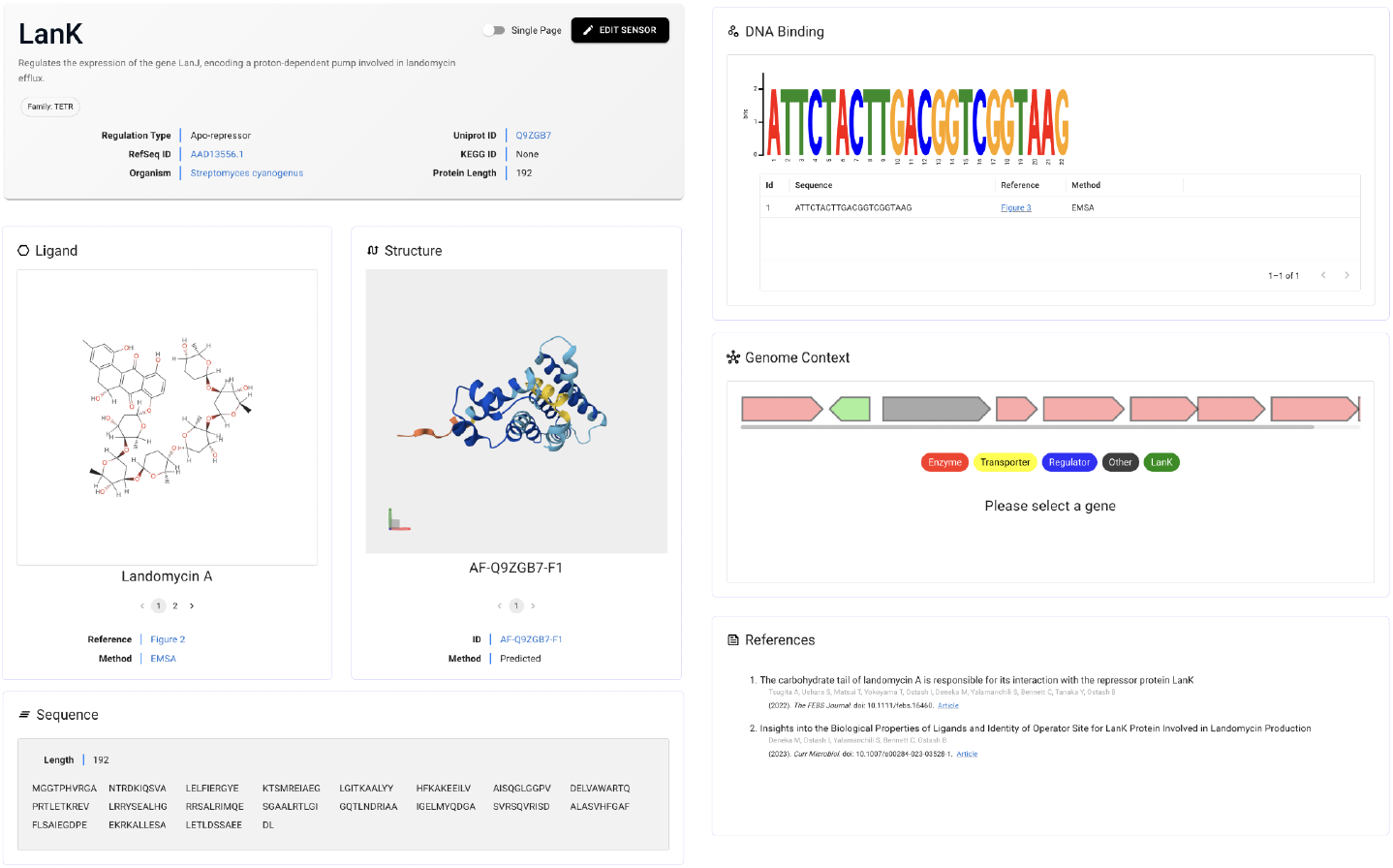
The user interface for transcription factor entry pages. Each page includes links to external databases, 2D structures of chemical inducers, interactive 3D protein structures, the protein sequence, DNA binding motifs, an interactive operon model, and associated references.

### Expanded content and statistics

The current release of groov^DB^ has more than doubled in size compared to its original release (**Table 1**). This version documents 588 manually curated binding interactions from 441 literature articles, which includes 320 unique ligand interactions and 268 unique DNA interactions. Data is distributed across 216 ligand-inducible TFs, which belong to over 8 major structural families, marking a substantial increase in database quality and quantity. There are now 450 unique crystal structures in groov^DB^ compared to 256 in the previous release. The TetR family is most represented, with over 100 members, likely due to their abundance in microbial genomes and their relative ease of characterization, since they are small, generally soluble, and often bind to short sequences (**Figure 2A**)^44^. The “Other” category of structural families includes 13 transcription factors belonging to the TrpR, ROK, DeoR, ArgR, EryD, and AsnC families. A wide range of structurally diverse ligands are represented, with two distinct clusters around sugars and coenzyme A ligated molecules (**Figure 2B**). Certain transcription factor families tend to bind to structurally similar ligands, such as the LacI family’s preference for sugars and the LuxR family’s preference for homoserine lactone quorum molecules. However, other families bind to many structurally unrelated molecules, including the TetR and MarR families.

**Table 1:** Comparison of the past and current versions of groov^DB^

| Content | groov<sup>DB</sup> 2022 | groov<sup>DB</sup> 2025 |
| --- | --- | --- |
| Sensors | 101 | 216 |
| Unique ligands | 131 | 320 |
| Unique organisms | 62 | 112 |
| DNA interactions | 101 | 268 |
| Entries with structures available | 34 | 216 |
| Unique crystal structures | 256 | 450 |

**Figure 2:**
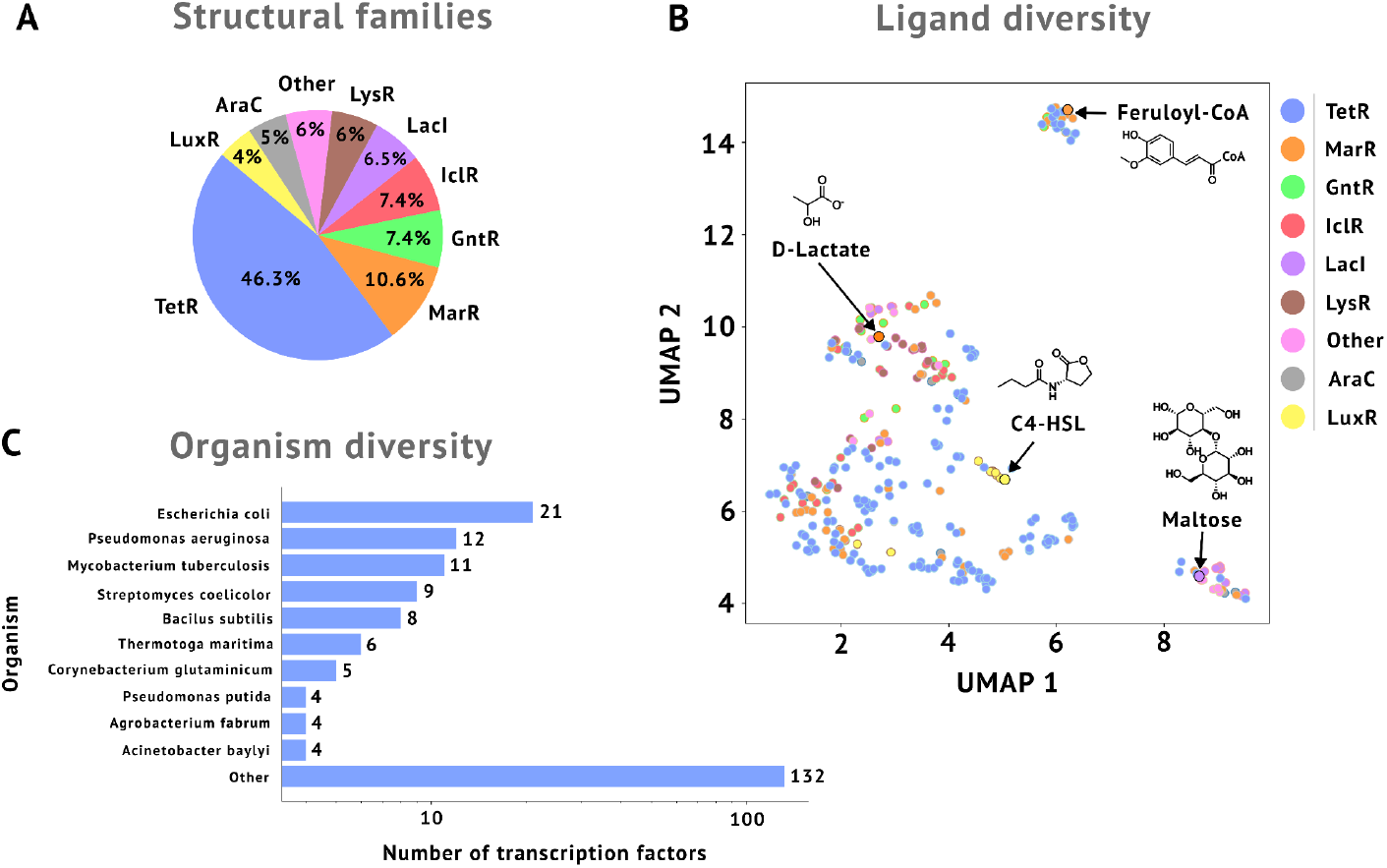
Content of groov^DB^ in 2025. (A) Relative abundance of major structural families. (B) UMAP projection of ligand chemical fingerprints colored by transcription factor family. Each point represents a unique ligand and selected ligands are highlighted and labeled. (C) Diversity of organisms represented in groov^DB^. Transcription factors belonging to the ten most represented organisms are highlighted.

Transcription factors within groov^DB^ belong to a total of 112 unique organisms, which are distributed among eight phyla, 11 classes, and 29 orders. Interestingly, although well-studied model organisms like *Escherichia coli* and *Pseudomonas aeruginosa* have greater representation, two thirds of all regulators come from organisms with four or fewer transcription factors, highlighting the diversity of organisms represented in groov^DB^ that would be missed in the more common organism-specific databases (**Figure 2C**). While groov^DB^ contains only a small subset of known transcription factors (21/304 in *E. coli*, 12/371 in *P. aeruginosa*, and 11/275 in *B. subtilis*) it is focused exclusively on ligand-inducible transcription factors with experimentally validated interactions to both ligands and DNA sequences^26,45,46^.

### Improved data infrastructure

In the previous version of groov^DB^, sensor pages would typically take one to three seconds to load, impeding user engagement. To provide a more rapid and robust infrastructure to host content in groov^DB^, we have implemented several significant changes to the architecture of the application. All database content is stored in two locations: a static JSON file for rapid access and a flexible NoSQL database for managing entry additions and updates. By fetching content from a static server, the web application is over five-fold more responsive compared to the previous version, based on the amount of time needed to load sensor pages. Both databases are always kept in sync with each other, which enables this hybrid database approach to strike a balance between speed and flexibility. To further improve responsiveness, we integrated the global React state manager, Zustand, which caches web page content. To provide a secure administration portal for supporting the integrity and accuracy of database content, we have deployed authentication endpoints using the Infrastructure as Code model. To ensure a robust framework that can scale with variable traffic, groov^DB^ was built using a serverless architecture that is able to expand and contract on demand. Finally, the frontend interface is deployed with AWS Amplify, which allows for a flexible continuous integration / continuous deployment workflow.

### Summary and future directions

The goal of groov^DB^ is to organize the world’s knowledge of genetically-encoded small molecule biosensors. The release of groov^DB^ presented herein represents a significant advance towards this mission by adding community-editing, enhanced visualizations, new query methods, and expanded content. In contrast to existing transcription factor databases, such as RegulonDB and RegPrecise, the modern software stack behind groov^DB^ aligns well with state-of-the-art practices that facilitate maintenance and future developments.

Beyond technical improvements, the expansion groov^DB^ content also provides scientific insights into the association between TF structural families and the ligands they bind. In agreement with observations from others, the TetR and MarR families bind a wide range of structurally diverse ligands, while the LuxR and LacI families tend to bind homoserine lactones or sugars, respectively^44,47–49^. As groov^DB^ grows, a higher resolution mapping between protein structure and inducer structure will likely emerge.

In the past couple of years we have directed our efforts towards making database contribution easier to promote the expansion of content. In the future, we plan to improve the data quality, quantity, and utility. Quality can be improved by supporting structural models of transcription factors in their multimeric forms, since most regulators in groov^DB^ form dimers. Content can be expanded by including TF interactions predicted via automated annotation, such as sequence homology or protein language models. Finally, the utility of groov^DB^ can be improved by tailoring it for practical use cases. We envision this involving the addition of more complex sequences, such as TF-regulated promoters and experimentally validated inducible expression plasmids, which can find immediate use within biotechnology workflows. We also aim to incorporate non-natural TFs, such as TF variants that have been engineered for improved sensitivity, that might be more practical for various bioengineering applications.

## Data availability

All original data containing experimental evidence displayed in groov^DB^ transcription factor entry pages are listed under the “References” section, with links to the source articles. All source code of groov^DB^ is freely accessible on GitHub under an MIT license. The GitHub repository for the frontend code is available here: https://github.com/groov-bio/groov-db-ui. The GitHub repository for the backend code is available here: https://github.com/groov-bio/groov-db-api.

## Acknowledgements

We are grateful for helpful suggestions from Khalid K. Alam, Adam J. Meyer, and R. C. Baer.

## Author contributions

J.D.L.: Led software development, and helped revise the original draft. M.T., B.M. R., N.Z., and P. T.: Data curation. M. S. and P.A.S.: Supervision and writing. S.D.: Conceptualization, software contributions, data curation, and writing.

